# Genetic mapping and nuclear interactions of an incompatibility response in *Agaricus bisporus*

**DOI:** 10.1101/2025.08.15.670591

**Authors:** K. Scholtmeijer, B. Auxier, Patrick Hendrickx, B Lavrijssen, A.J.M. Debets, JJP. Baars, DK Aanen, A. van Peer

**Author notes:** These authors had equal contributions to this manuscript (K. Scholtmeijer, B. Auxier).

## Abstract

During cultivation, mixing of different heterokaryotic individuals of the button mushroom, *Agaricus bisporus*, generally reduces yield. This phenomenon could be caused by direct antagonistic responses and/or reduced synchronization by not forming a chimeric hyphal network. In other fungi, highly divergent alleles for a set of genes affect successful network formation between individuals either by preventing fusion or, more commonly, triggering cell death post-fusion. To understand this process in *A. bisporus*, it is important to identify the allelic variants allowing these fungi to discriminate self from nonself. We leverage a recently described cell death staining method utilizing Evans Blue to visualize mycelial compatibility. Here, we provide results of a first genetic mapping of incompatibility alleles in *A. bisporus*. Crossing strains between *A. bisporus* var. *bisporus* and *A. bisporus* var. *burnetti* we find segregation ratios of compatible progeny generally consistent with three nuclear loci. To identify these regions, we first use a set of single Chromosome Substitution Lines (CSLs), produced by genotyping progeny with recombination skewed to the very chromosome ends. We localize the main effect to be between two and three chromosomes, depending on the common nucleus of interacting heterokaryons. Using genome-wide markers for 167 sexual progeny, we identify loci controlling mycelial compatibility on chromosomes 4, 6 and 7, the same chromosomes as indicated by chromosome substitution lines. Notably, while the choice of a common nucleus seemed to affect the compatibility of CSLs, it did not seem to affect the loci identified in the sexual progeny. The ability to mix different strains of this mushroom-forming fungus could allow additional cultivation approaches, combining strains with complementary characteristics. These results provide a starting point towards understanding the molecular mechanisms underlying this fundamental property of hyphal networks in basidiomycetes.

## Introduction

The distinction between self and nonself interactions in fungi is fundamentally important. As the thread-like hyphae of an individual, the mycelium, grow through a substrate, they can encounter hyphae of other individuals. When this occurs between hyphae of the same species, generally the two hyphae fuse, and if they are from different individuals this fused cell is rapidly killed (Garnjobst & Wilson, 1956). This prevention of fusion between nonself individuals has the benefit of preventing the spread of deleterious elements like nuclei or viruses (Debets & Griffiths, 1998; Grum-Grzhimaylo et al., 2021). In Basidiomycetes, the distinctive heterokaryotic state, with two distinct haploid genotypes in a common cytoplasm, presents additional vulnerabilities as mating must accommodate the fusion between two homokaryotic individuals (Auxier et al., 2022; James, 2015; Rayner, 1991). This incompatibility is expressed as two potentially overlapping phenomena; *mycelial compatibility* where secondary characteristics of mycelia such as metabolite secretion or aerial hyphae produce a zone of interaction aside from hyphal fusion, and *somatic incompatibility* where the fusion of hyphae between two individuals leads to cell death in the fusion zone (Worrall, 1997).

This cell death is caused by cytoplasmic proteins, whose interaction presumably triggers downstream processes causing death of the fused cell (Saupe et al., 1996). For sustained fusion between two hyphae, the two cytoplasms must encode the same versions of the respective proteins. This is a version of synthetic lethality, where cell death is triggered by a combination of alleles not seen in nature (Dobzhansky, 1946). For sustained fusion between two hyphae, the two cytoplasms must encode the same versions of the respective proteins. The alleles encoding these proteins form a sort of barcode that must match to allow stable fusion (Gonçalves & Glass, 2020). Sustained fusions between hyphae allows for persistent social interactions, functionally defining an individual. The molecular details of this system have been worked out in some detail in Ascomycete fungi, particularly *Neurospora crassa* (Daskalov et al., 2020) and *Podospora anserina* (Clavé et al., 2024) and between 10 and 15 genes each with multiple alleles have been identified. This high number of genes makes it unlikely that unrelated individuals have the same alleles for all loci, thus generally preventing fusion between different individuals and providing an effective non-self recognition system (Gonçalves & Glass, 2020; Nauta & Hoekstra, 1994). However, the genes responsible for this in Basidiomycetes are unknown.

Previous studies on the genetics of incompatibility in several mushroom-forming species indicates that generally several loci are involved. In *Amylostereum* it was indicated that at least two loci segregated between two individuals (van der Nest et al., 2008), and in *Heterobasidion annosum* three to four loci (Hansen et al., 1993; Lind et al., 2007). The wood-rotting *Serpula lacrymans* had three loci segregating, although this species seems to have low genetic diversity overall (Kauserud et al., 2004, 2006). In the wood-decay fungus *Phellinus gilvus*, it was found that a single locus was the primary contributor to the phenotype (Rizzo et al., 1995). Those studies suggest that the number of loci regulating incompatibility in Basidiomycetes is lower than in Ascomycetes.

Currently, button mushroom cultivation requires the use of single strains. This is because cultivation of mixed cultures leads to dramatic reductions in yield (Kerrigan & Wach, 2014; Scholtmeijer et al., 2025). This incompatibility response between heterokaryons has also been demonstrated to prevent virus transmission (O’Connor et al., 2021). In theory, knowing the genes, or at least narrow genetic regions, causing incompatibility could allow selection of individuals that would be somatically compatible. Availability of somatically compatible, but genetically diverse, strains could allow selection on more specialized aspects of cultivation.

Here, we present results from mapping the incompatibility loci in *A. bisporus*. Using a set of single chromosome substitution lines as well as sexual *A. bisporus* var. *bisporus* X *A. bisporus* var. *burnetti* populations of meiotic progeny, we demonstrate the segregation of three loci. These loci were genetically mapped but could not be resolved to a gene level. To understand the interaction between nuclei in a heterokaryon, we leveraged the multiple mating alleles to mate the progeny with different mating partners. Comparisons between these populations showed that the loci segregating seemed to be independent of the partner nucleus, indicating dominant interactions.

## Materials and Methods

### Terminology

For clarity, here we use ***heterokayon*** to refer to a fungal individual with two nuclear genotypes, necessary to produce mushrooms. We use ***homokaryon*** to refer to isolates with a single type of nuclei, which can be derived either from screening the often heterokaryotic sexual spores for rare homokaryotic ones, or from protoplasting to isolate individual genotypes. Two homokaryons can be ***mated*** to produce a heterokaryon, assuming the mating-type loci are compatible, and if mushrooms are produced leading to sexual spores, we refer to this as ***crossing***. Following the terminology of Worrall, we use ***common nucleus*** when we independently mate different homokaryons with the same common homokaryon, producing related heterokaryons. Finally, we reserve the term ***interaction*** for testing incompatibility between heterokaryons. In this work, matings occur exclusively between homokaryons, and interactions are exclusively tested between heterokaryons.

### Strains and Culture conditions

Our starting material was two heterokaryotic *A. bisporus* strains, two-spored A15 and four-spored Bisp015 (Table 1). We recovered the two homokaryotic genotypes of each heterokaryon by protoplasting (Sonnenberg et al., 1988), using PCR with mating locus allele specific primers to identify the two component nuclei. The two recovered homokaryotic components of A15 we designated as A and B, and the two from Bisp015 (Kerrigan, 1995, 1996) as C and D. These strains have compatible mating types in all six pairwise combinations. To produce CSLs (see below), we used two homokaryons, H39 (clonally related to A) and H97 (clonally related to B). The strains listed and genetic codes are found in Table 1. Note that we hereafter refer to E as A and F as B, due to the clonal relationship. For routine culture, cultures were grown on MMP Agar (Scholtmeijer et al., 2025) at 24 °C.

**Table 1:**
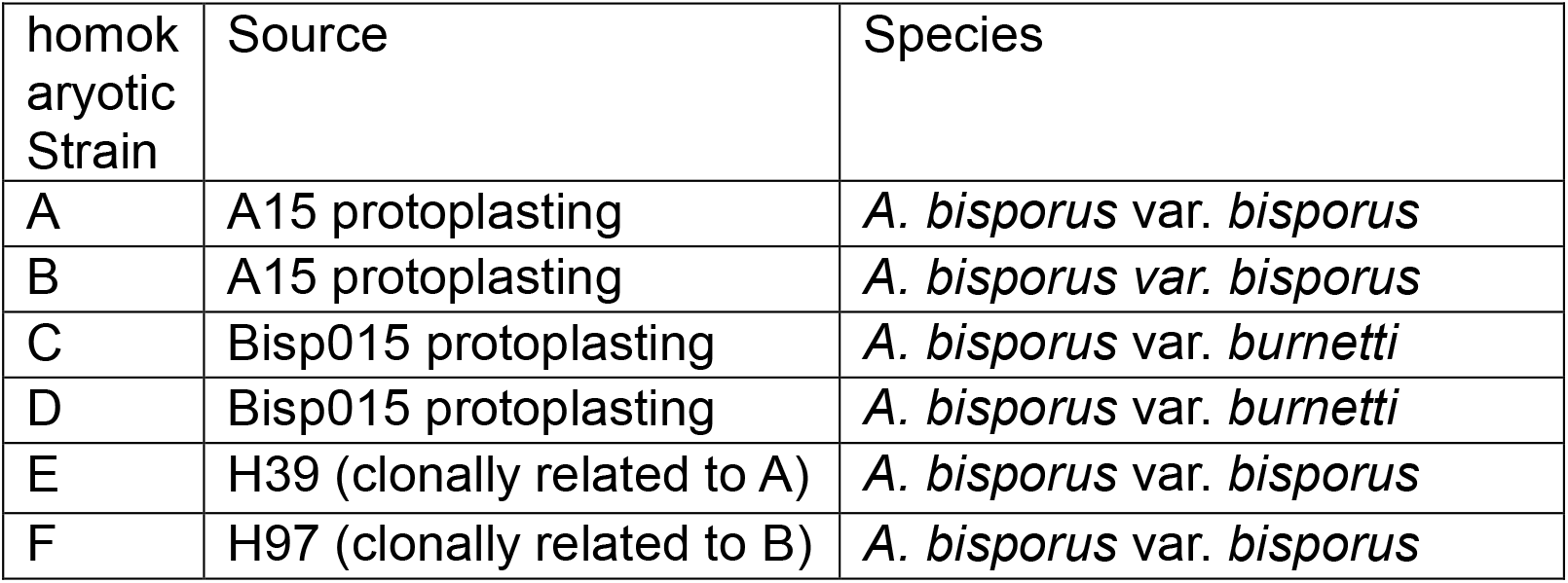
*A. bisporus* strains, all with compatible mating types, used in genetic crosses

### Chromosome Substitution Lines

To identify which chromosome(s) harbor alleles for the incompatibility interaction, we produced a set of single chromosome substitutions lines (CSLs). For this, we crossed E and F, and screened basidiospores for colonies with a single mating type, i.e. homokaryons (see below). Using 42 SNP markers spread across the chromosomes, we genotyped the single-spore isolates (SSI) to recover isolates with essentially non-recombined chromosomes, which is possible as recombination in *A. bisporus* var. *bisporus* predominantly occurs at the chromosome ends (Sonnenberg et al., 2016) and with the genetic background of H39 except for one of the thirteen chromosomes of H97 (Figure 1A). We produced one of each CSL except for chromosomes 4 and 7, for which we recovered two independent CSLs.

**Figure 1:**
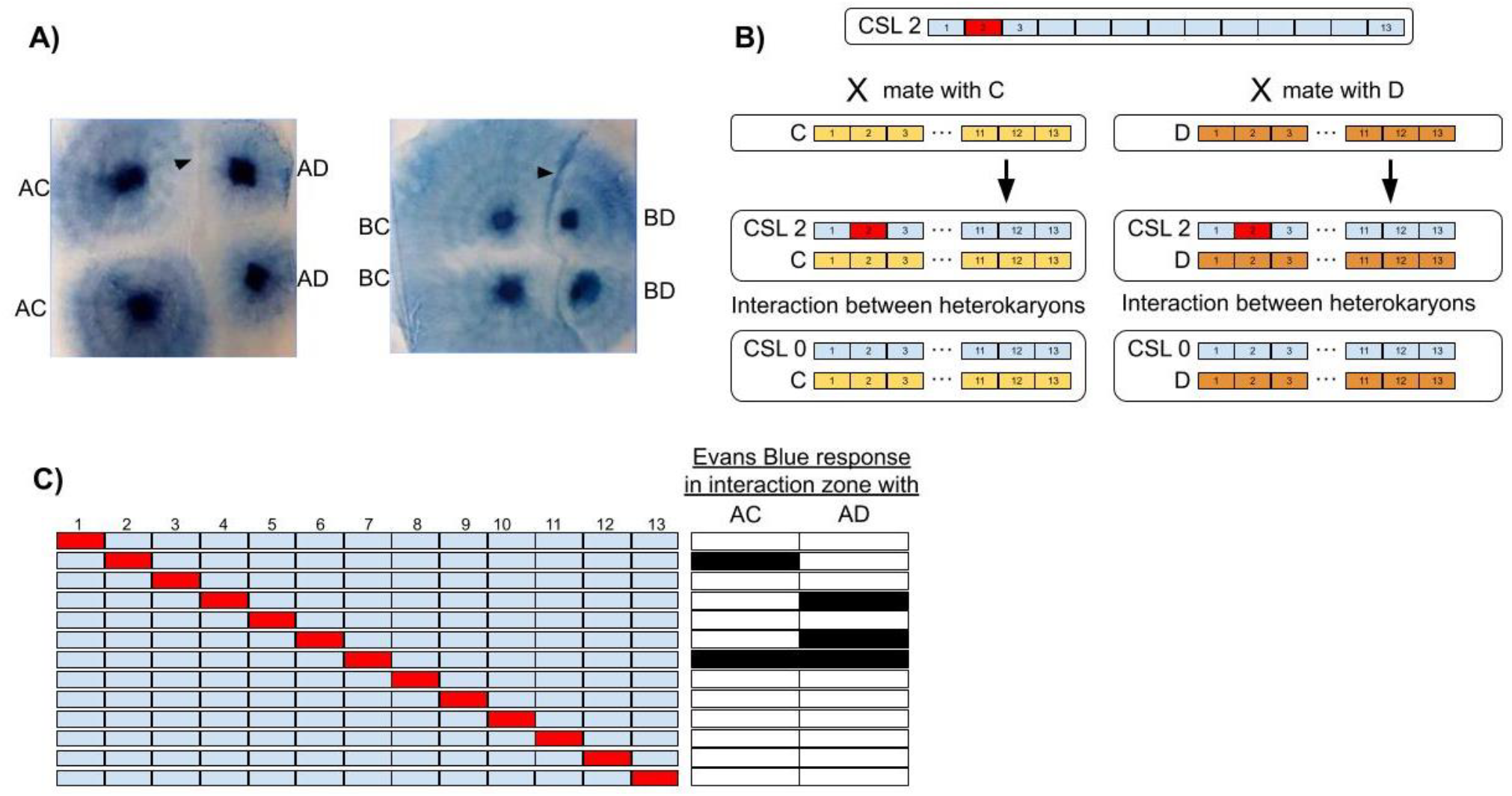
Incompatibility assessed between *A. bisporus* strains A and B, when mated with C or D isolates. A) incompatible response under Evans Blue staining, with a dark blue line forming between the different heterokaryons (arrowheads) but not between self-interaction controls. B) shows the construction of the CSLs and subsequent interaction testing. C) Results of interactions scoring for the 15 CSLs tested against AC or AD heterokaryons. Black boxes indicate incompatible responses due to the corresponding B chromosome in the CSL.

These homokaryotic CSL isolates were then mated with either C or D homokaryons, and interactions with AC or AD were measured using Evans Blue staining as described previously (Scholtmeijer et al., 2025). We use this as a measure of incompatibility, as this stain is excluded from living tissue and thus highlights dead hyphae.

### Meiotic progeny

To produce recombinant meiotic progeny, we crossed B and D, collecting single spore isolates (SSI) from spore prints from multiple mushrooms. These SSI were screened for homokaryotic progeny using primers designed for the specific allele of the HD mating-type locus (Tables S1 and S2). These meiotic progeny were mated with either A or C, and incompatibility interactions were also scored as for the CSLs. In addition, we produced a second meiotic population by crossing A with C. These were treated as above, but mated instead with either B or D.

### Genetic mapping

For the population resulting from the cross of B with D, we genotyped in two differing ways. First, we genotyped 96 homokaryotic SSI with 106 SNP markers using a competitive allele specific PCR (KASP assay). To this population, we added 71 additional isolates, which were processed for whole genome sequencing with paired-end 150 bp short reads. Reads were aligned to the *A. bisporus* H97 version 3.1 reference genome (Sonnenberg et al., 2016) with bwa-mem v.0.7.17 (Li, 2013) using the default settings and variants were called with Freebayes v1.1.0 (Garrison & Marth, 2012) using default settings. The resulting variant file was filtered to retain only variants of type SNP with a minimum read depth of 10, and the filtered variant file was converted in a genotype file using dedicated scripts. The genotype at the same 106 SNP loci was determined and merged into one genotype file.

Genetic mapping was performed using the R/qtl2 package (Broman et al., 2019). We removed any individuals with >25 markers with missing genotypes and removed potential clones. We removed one marker, Ch5_6859, that showed segregation distortion (0.86 frequency). After filtering, the dataset consisted of 96 markers and 147 individuals. We constructed a linkage map using the minimum spanning tree approach implemented in ASMap (Taylor & Butler, 2017; Wu et al., 2008), resulting in 23 linkage groups using a 1×10^−4^ significance threshold. We manually merged the 23 linkage groups into the 13 described chromosomes (Sonnenberg et al., 2020). We visualized the resulting genetic map with LinkageMapView (Ouellette et al., 2018).

We mapped the incompatibility trait using Haley-Knott regression using the binary model as our phenotypes are binary. We determined significance thresholds using 1000 permutations of the data, using α= 0.05. Significantly associated windows of the genome were based on the peak LOD value bordered by a drop of 1 in LOD from this peak on either side.

## Results

### Genetic basis for incompatibility between *A. bisporus* isolates

Initial tests showed that heterokaryons we produced displayed an incompatible response, of variable strength, between isolates (Figure S1). This incompatible response was not observed between self-pairing of isolates (Figure S1). For example, when we mated A with C and B with C, interactions between this AC heterokaryon and the BC heterokaryon produced increased Evans Blue staining in the interaction zone (Figure 1B, arrowhead), while self-interactions of AC with AC or BC with BC did not produce an incompatible response. Similarly, A or B were mated with the common nucleus D, the interactions between AD and BD heterokaryons produced an incompatible response, while self-interactions did not (Figure 1B).

To determine the chromosomes associated with this response, we produced a set of single chromosome substitution lines (CSLs) (Figure 1A), a set of lines with twelve chromosomes of A and a single non-recombinant chromosome from B. Genotyping across 42 markers, we identified strains that were essentially the genotype of A, but with one B chromosome (Supplementary File 1). We recovered a single CSL for each chromosome, with duplicates for chromosome 3 and 4. By mating these 15 CSLs with either homokaryon C or D, we produced two sets of heterokaryons. These two sets of heterokaryons were tested for incompatibility against the AC heterokaryon (common nucleus C) or AD (common nucleus D) using Evans Blue staining to detect cell death in the interactions zone (Figure S2). When mated to the common nucleus C, among the set of CSLs those with Chromosome 2 and 7 from B showed incompatibility to AC (Figure 1C; Figure S2). When mated with common nucleus D, CSL-4,6, and 7 showed incompatibility with AD (Figure 1C; Figure S3). For the duplicate CSLs for Chromosome 3 and 4, the results were consistent (Figure S3). Chromosome 1, which contains the mating type locus, showed no involvement in incompatibility.

### Segregation of incompatibility within recombinant populations

From the B/D population, we attempted to mate all 92 progeny with both A and C. When mated with strain A, 14.3% of B/D offspring were compatible with AB, and 9.4% with AD. We observed that 16 individuals (10.9%) did not successfully mate with strain A. When we instead mated the 92 progeny with isolate C, we found that 14.6% were compatible with the BC heterokaryon, while only 5.4% were compatible with the CD heterokaryon.

We also crossed A with D, collecting 98 homokaryotic SSIs from the A/D population, which were mated with either strain B or C. When mated with B, 11% of this A/D population was compatible in interactions with an AB dikaryon (Table 2; top row). Likewise,12% of the A/D population was, when mated with C, compatible with the CD heterokaryon (Table 2; fourth row). Both values were statistically consistent with three locus segregation, with a p value >0.05 from an exact binomial test.

**Table 2:** Segregation of compatible reactions in hybrid progeny pools

| Set* | Pop. | Mating Partner | Interact. individ. | Interaction notation<sup>1</sup> | Segregation testing | Compatible Frequency<sup>2</sup> | Three loci hypothesis<sup>3</sup> |
| --- | --- | --- | --- | --- | --- | --- | --- |
|  | (A/D) | B | AB | (A/D + <u>B</u>) vs. (A + <u>B</u>) | D different from A | 11% (98) | p=0.87 |
|  |  |  | BD | N/D | N/D | N/D | N/D |
|  |  | C | AC | N/D | N/D | N/D | N/D |
|  |  |  | CD | (A/D + <u>C</u>) vs. (<u>C</u> + D) | A different from D | 12% (98) | p=1.00 |
| 1 | (B/D) | A | AB | (B/D + <u>A</u>) vs. (<u>A</u> + B) | D different from B | 14.3% (84) | p=0.61 |
| 2 |  |  | AD | (B/D + <u>A</u>) vs. (<u>A</u> + D) | B different from D | 9.4% (75) | p=0.48 |
| 3 |  | C | BC | (<u>B</u>/D + <u>C</u>) vs. (B + <u>C</u>) | D different from B | 14.6% (82) | p=0.50 |
| 4 |  |  | CD | (B/D + <u>C</u>) vs. (<u>C</u> + D) | B different from D | 5.4% (92) | p=0.04 |

\* used for QTL mapping in Figure 4
<sup>1</sup> underlined letters indicate shared genotypes between the two heterokaryons, note N/D = Not Done
<sup>2</sup> numbers in brackets indicate total number of progeny that were used for phenotyping
<sup>3</sup> p values represent exact binomial test assuming segregation of compatible progeny at 0.125.

### Genetic map of the *A. bisporus* var. *bisporus* X *A. bisporus* var. *burnetti* BD cross

To increase resolution of genetic mapping, we crossed B with genotype D, a homokaryon from the closely related *A. bisporus* var. *burnetti*, a cross described to have an increased frequency of non-telomeric crossovers (Foulongne-Oriol et al., 2010). Sequencing of these meiotic progeny showed a total genetic distance of 1326.8 cM spread across the 13 chromosomes (Figure 2). Particularly for the larger chromosomes, despite even large physical distances between markers, genetic distances were highly skewed, with large portions of chromosomes 2, 3, 4 and 7 being essentially non-recombinant.

**Figure 2:**
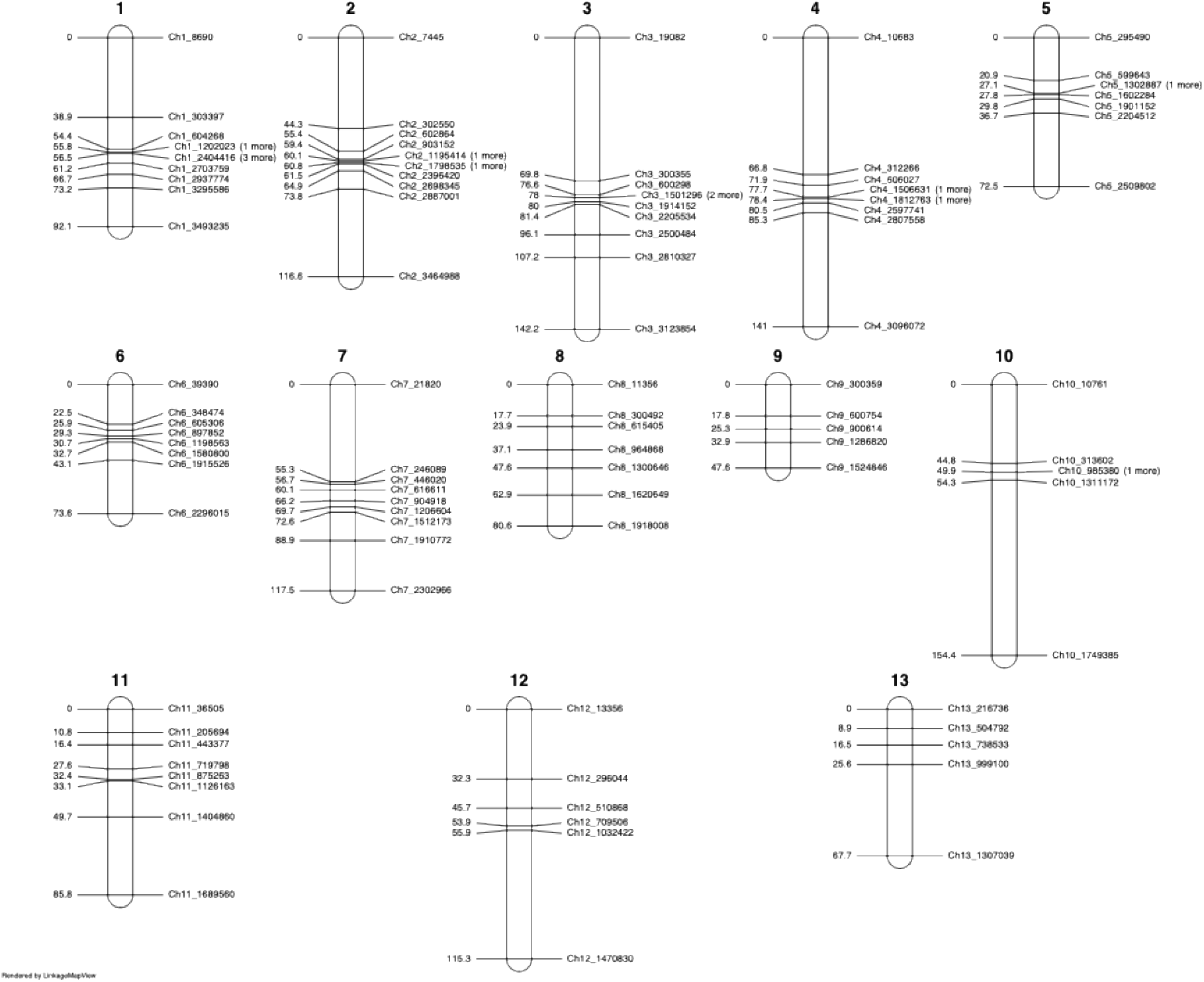
Genetic map of meiotic progeny of A (*A. bisporus* var. *bisporus*) X D (*A. bisporus* var. *burnetti*). Spacing of the 108 markers across the 13 chromosomes is depicted, scaled along the chromosomes based on cM distance (left side). Physical position is found in the marker name, with markers being spaced approximately 0.3Mb apart.

### Mapping of incompatibility within the (B/D) population

The Evans Blue phenotype across the offspring was variable, with some isolates producing less staining overall (compare left and right isolates in Figure S4). Compatible interactions generally had less staining than the mycelia near the inoculation site (Figure S4 A-D). When incompatibility was observed, as a dark blue line between the genotypes, it varied between a weak interaction zone (Figure S4 E-F), to a darkly pigmented zone (Figure S4 G-H). For the subsequent genetic mapping, we split the interactions into either compatible or incompatible, lumping all gradients of weakly/strongly incompatible into two categories.

We phenotyped the interaction of the B/D population against four different heterokaryons (Table 2). Using genetic information from 96 markers spread across the 13 chromosomes, we mapped the loci allowing compatibility. QTL mapping results in peaks on chromosome 4, 6 and 7 being found in the AB, AD, and BC interactions (Figure 3). A significant peak was also found on chromosome 10 for interactions involving the AB individuals. Interactions involving the CD individual did not result in any significant peaks.

**Figure 3:**
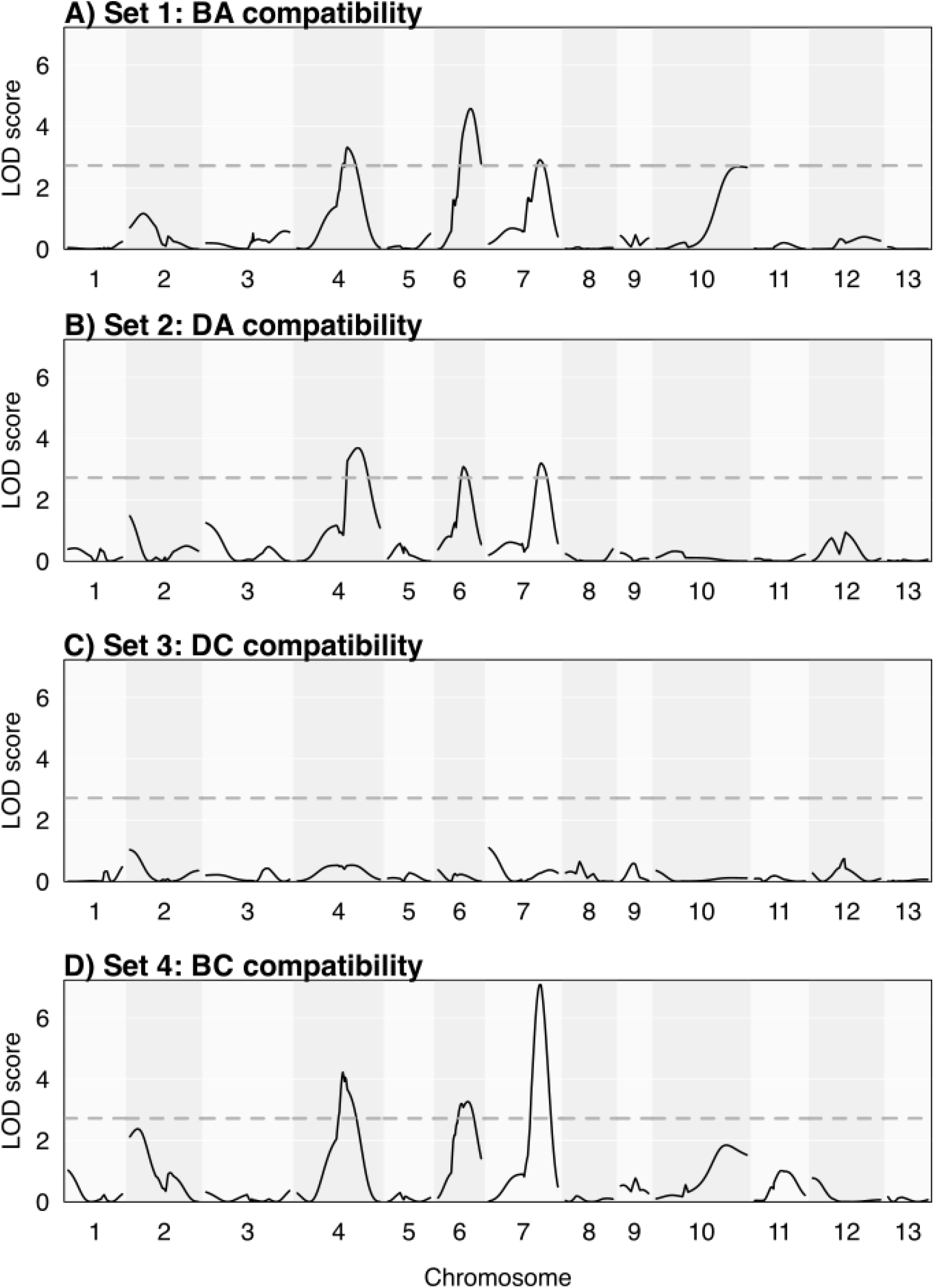
QTL mapping of compatibility within the B/D population. Panels A-B show interactions of B/D progeny when mated with A and interacted against either BA or DA heterokaryon. Panels C-D show results when the same population was mated with C and interacted with DC or BC. Different chromosomes are indicated by differential vertical shading. Statistical significance is indicated by horizontal dotted line, and was determined by a permutation test.

## Discussion

Our results show that several loci contribute to the incompatibility phenotype of *A. bisporus,* spread across multiple chromosomes. In the meiotic progeny, genetic mapping showed a consistent association on chromosomes 4, 6 and 7. Across the 4 common nuclear backgrounds tested, we recovered peaks centered at similar regions of the chromosomes. This likely indicates that the effect of the common nucleus is not major in this experiment. In interactions between offspring of B and D when mated with C, the compatibility ratio compared to DC segregated significantly different from a three-locus model, although such a marginally significant result is not unexpected when testing across multiple backgrounds.

Our use of CSLs allowed for rapid screening of the incompatibility phenotype, and likely would be of use for screening other dominant phenotypes. Such segregating populations are used in species like *Aspergillus niger*, as well as plants like *Arabidopsis thaliana* among others (Debets et al., 1993; Wijnen et al., 2024). One caveat of the lines used here is that the CSLs are not composed of entire non-recombinant chromosomes, as there is telomeric recombination. While it is possible that some of the nonself recognition genes are located near telomeres, as seen in *Neurospora crassa* (Wang et al., 2023), this is not likely in our experiment. The meiotic crossing data showed that the identified QTL were localized to the center of the chromosomes.

In the model Ascomycete, *Neurospora crassa*, it was noted early on that the mating type was involved in nonself recognition (Garnjobst, 1953). The molecular details of this interaction have been determined (Jacobson, 1992; Shiu & Glass, 1999), and it is often thought that this is a common factor in fungal nonself recognition. Here, we do not find any association between the chromosome where the functional mating locus resides, chromosome 1, and incompatibility. This lack of association may be more common, as even in most Ascomycetes it is not found (Auxier et al., 2024; Choi et al., 2012). These data are consistent with our previous model, where compatible mating types allow “escape” from the incompatibility response due to rapid nuclear migration, and not transcriptional regulation (Auxier et al., 2021).

It has been suggested previously that using an experimental design where a genotype is shared across heterokaryons, a so-called common nucleus, can simplify the genetic analysis (Worrall, 1997). Our experiments indicate this may not be a safe assumption, as the comparison between sets 2 and 4 show significantly different segregation ratios. We observed that when paired with nucleus A many more offspring are compatible with D than when paired with nucleus C. It may be that the influence of the common nucleus is shielding the effects of an incompatibility locus, something we have speculated on previously (Auxier et al., 2021). Previous studies have made use of a common nucleus when mapping this trait (Hansen et al., 1993), but few have tested the effect of different common nuclei.

Combining the evidence of CSLs and meiotic progeny, we predict strong incompatibility alleles on chromosomes 4, 6 and 7. Finding three loci segregating between two haploid strains is consistent with findings from other Basidiomycetes like *Heterobasidion annosum* (Lind et al., 2007). We have also shown that segregation of the nonself recognition response was consistent with three loci in *Coprinopsis cinerea*, and that one locus contains components resembling nucleotide binding leucine-rich repeats genes (Auxier et al. *in prep*). However, whatever the ultimate mechanism, it is important to recognize this is significantly fewer than the number of loci that segregate in Ascomycete species (Gonçalves & Glass, 2020). As we have suggested previously, this may be due to the combination of two genomes inside a heterokaryotic mycelium (Auxier et al., 2021). If the alleles for these loci act co-dominantly, then a few loci could provide a large number of potential vegetative compatibility groups.

Bioinformatic methods to identify nonself recognition genes in Basidiomycetes have produced mixed results. Since many details of the molecular mechanisms of nonself recognition have been worked out in Ascomycetes, and genomic data is rapidly accumulating, large-scale surveys have attempted to transfer this knowledge to mushroom-forming fungi. An initial survey found Basidiomycete homologs for many of the genes used for this process in Ascomycetes (van der Nest et al., 2014). More detailed surveys of the NLR-based mechanisms also found many copies in Basidiomycete fungi (Dyrka et al., 2014). Recent updates in both genomic data and known mechanisms have increased the number of genes found, and highlighted that Basidiomycetes have particularly diverse NLR genes that require additional functional validation (Wojciechowski et al., 2022). However, these studies are limited by the fact that many NLR genes in Ascomycetes are involved in processes other than nonself recognition (Uehling et al., 2017), and so sequence homology has limited power to infer function.

The commercial cultivation of *Agaricus bisporus* means that somatic incompatibility is particularly relevant. Mixing of different genotypes can lead to dramatic yield reductions (Scholtmeijer et al., 2025). In effect, this adds a restriction to cultivation, since all members of the production chain must agree on a single clone to use. A more complete understanding of the incompatibility response may allow us to bypass this requirement. Additionally, one of the major diseases encountered in mushroom cultivation is Mushroom Virus X, which can drastically reduce yields (Ross et al., 1987). This virus consists of a cluster of ss(+)RNAs and is transmitted through hyphal fusion (Gandy, 1960; Grogan et al., 2003; O’Connor et al., 2021), and the incompatible response between individuals presumably limits virus transfer, although in laboratory conditions this can be broken through (O’Connor et al., 2021). The mechanism by which viruses can spread despite this cell death is unclear, but may be related to idiosyncratic interactions between virus types and specific incompatibility loci (Choi et al., 2012). With identification of these three loci, even in the absence of molecular characterization, it would be possible to test the contribution of these loci in controlling virus transmission.

## Conclusion

This work confirms that there is a genetic basis for incompatibility in *A. bisporus*, with a similar number of loci compared to other Basidiomycetes. Given the importance of somatic incompatibility at a basic biological level, as well as the industrial relevance, understanding this trait is an important future goal. While the specific genes underlying this trait remain unknown in this species, this work provides a path towards unraveling this genetic trait.

## Data Availability

All genotype data and code for analyses are available in the publicly available repository https://github.com/BenAuxier/Agaricus_mapping. Recombinant progeny are available from AvP upon request.

## Acknowledgements

The authors thank Chris Dijkstra for helpful comments.

## Funding

This project received funding from the Netherlands Research Organization (NWO) ALGR.2017.010

## Conflicts of Interest

JB is partially employed by CNC Grondstoffen. Neither this employer, or any other funding source reviewed the manuscript prior to submission. The authors have no other conflicts to declare.

## Supplemental Materials

**Figure S1:**
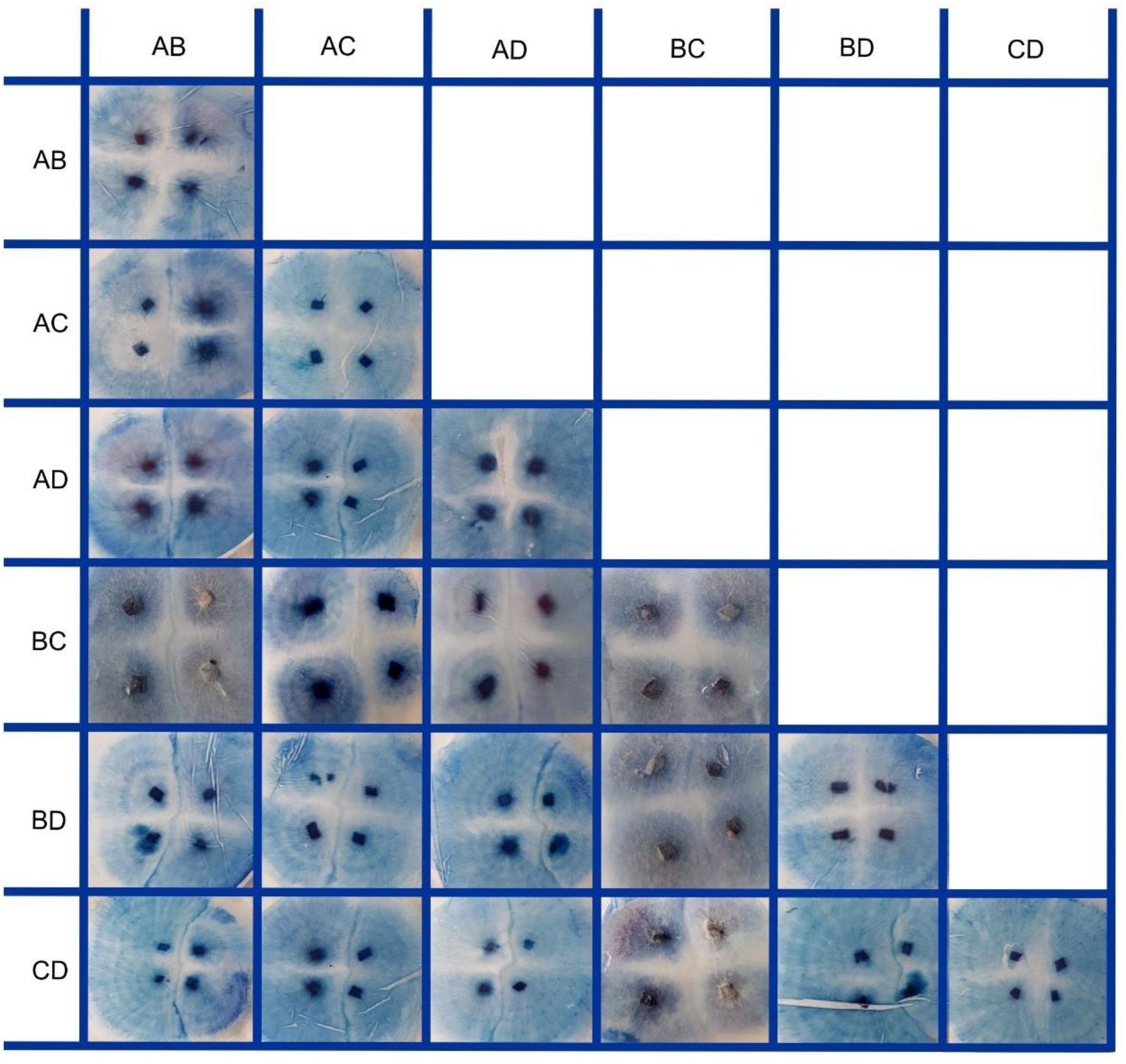
Interactions of heterokaryons synthesized from the four starting homokaryons. Note lack of interacting blue zone in self pairings (diagonal) as well as in off-diagonal pictures in vertical pairings (self). Blue lines result from cell death in pairings in all nonself interactions, except AC X BC and BC X CD, where the horizontal interactions produce little staining.

**Figure S2:**
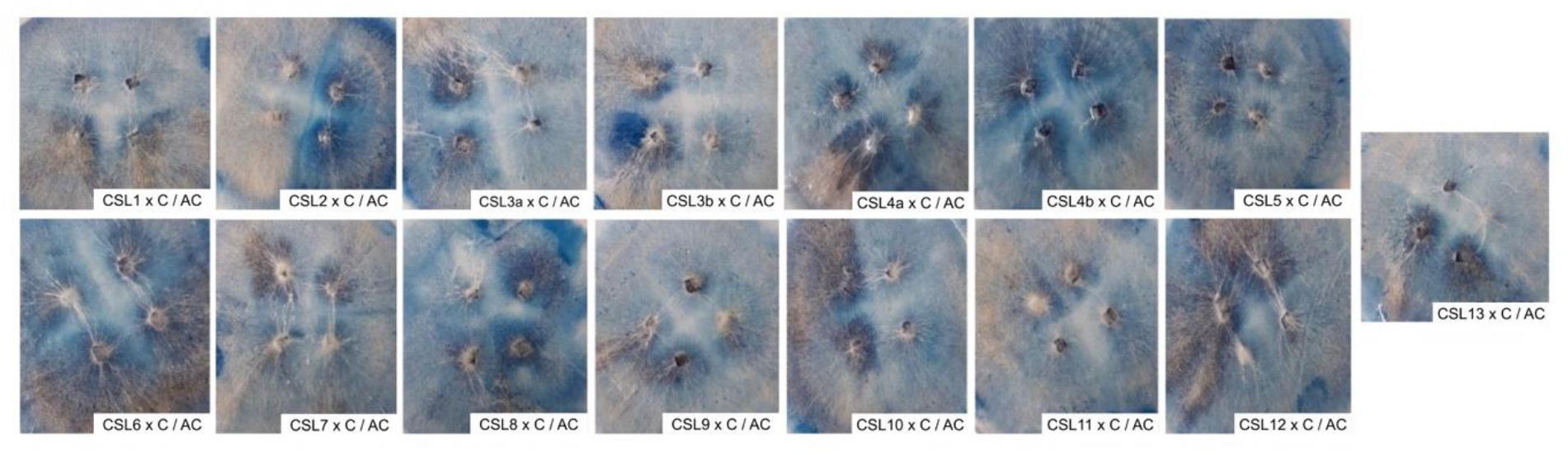
Interactions between CSLs mated with C and AC heterokaryon. Blue lines indicate CSLs for chromosome 2 and 7 are incompatible.

**Figure S3:**
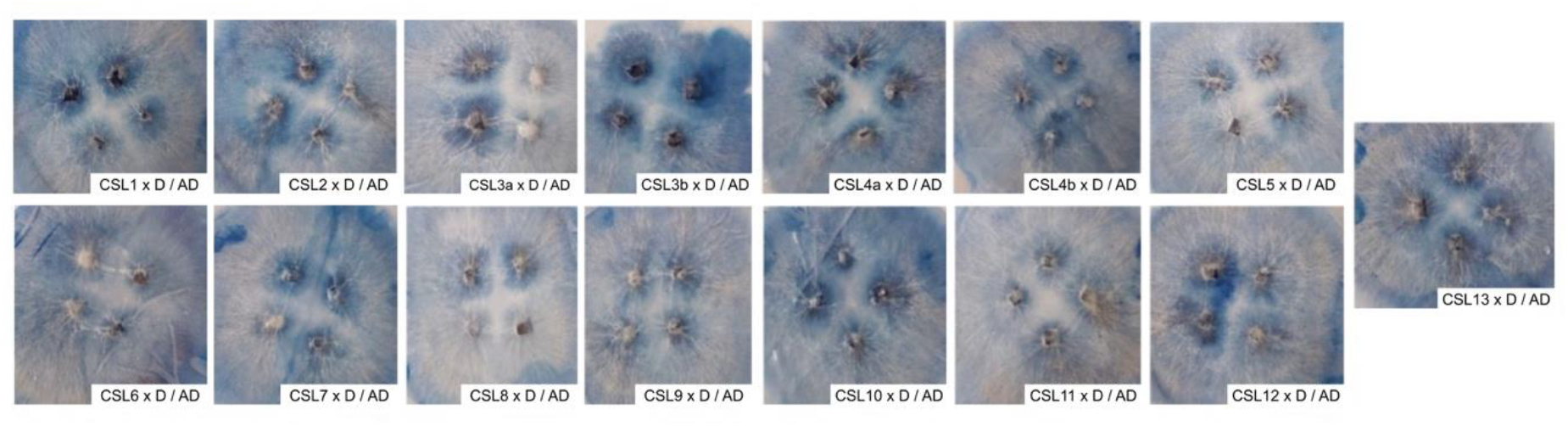
Interactions between CSLs mated with D and AD heterokaryon: CSLs for chromosome 4, 6 7 are incompatible with AD.

**Figure S4:**
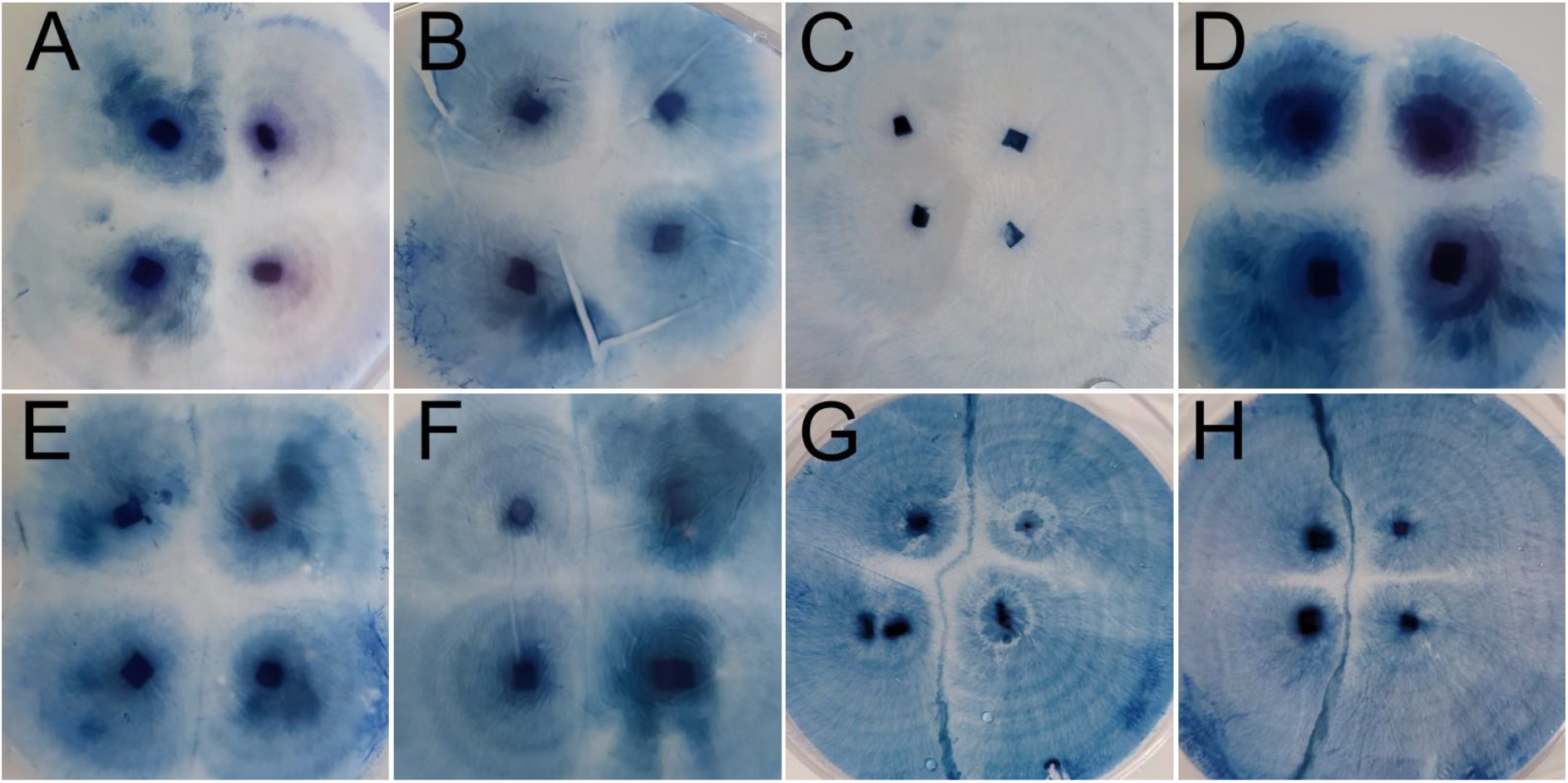
Representative phenotypes of incompatibility reaction. Interactions were setup as in Figure 1, with replicate individuals in the top and bottom rows, and different individuals on the left or right. Variation in interactions between meiotic progeny scored as compatible are shown in A-D, while weakly incompatible interactions are shown in E and F, and strongly incompatible interactions in G and H.

**Table S1:**
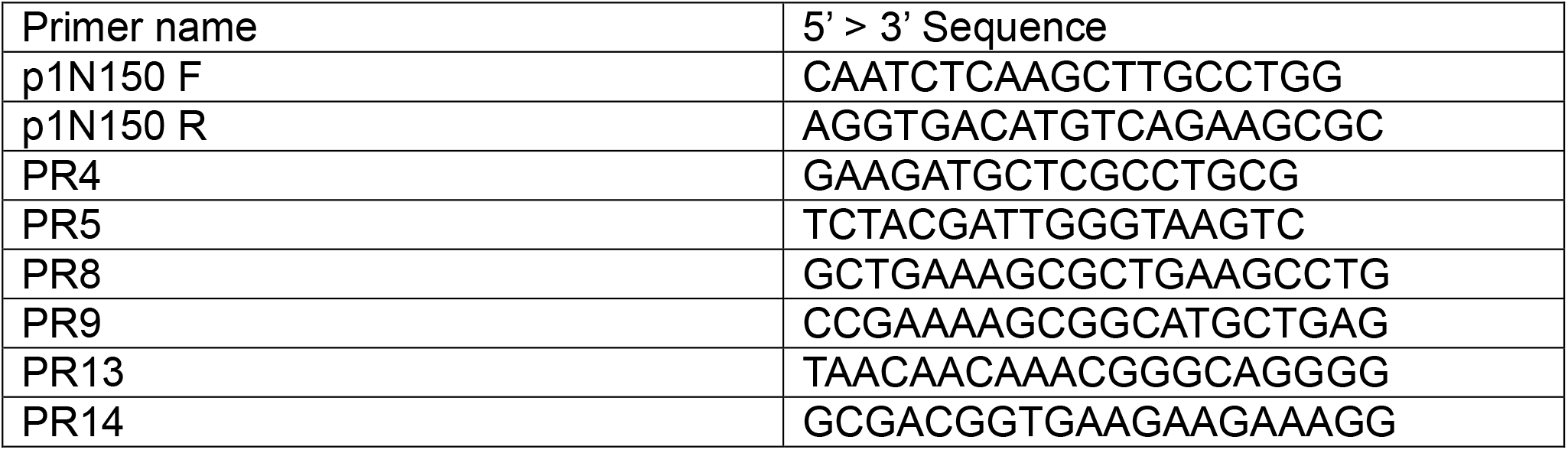
Primers used to identify mating types of isolates

**Table S2:** Predicted amplicon sizes for primer pairs from different mating types^1^

| Forward Primer | Reverse Primer | A15_p1 (A) | A15_p2 (B) | Bisp015_p1 (C) | Bisp015_p2 (D) |
| --- | --- | --- | --- | --- | --- |
| P1N150F | P1N150R | 750 bp | 1300 bp | 750 bp | 750 bp |
| PR4 | PR5 | 781 bp | - | x | x |
| PR8 | PR9 | x | - | 1111 bp | x |
| PR13 | PR14 | x | - | x | 1510 bp |

<sup>1</sup> Sizes indicated in base pairs, with “x” indicating no predicted product, and “-“ indicating untested combinations.

**Supplementary file 1**: Genotypes determined by KASP markers for the 15 single CSLs used.

